# The temporal regulation inter-leaves from domesticated-tomato contrasts with timelessness of its wild ancestors

**DOI:** 10.1101/2022.10.25.513690

**Authors:** João Antonio Siqueira, Auxiliadora O. Martins, Thiago Wakin, Marcelle Ferreira Silva, Willian Batista-Silva, Fred A.L. Brito, Alisdair R. Fernie, Adriano Nunes-Nesi, Wagner L. Araújo

**Affiliations:** Departamento de Biologia Vegetal, Universidade Federal de Viçosa, 36570-900 Viçosa, MG, Brazil; Max-Planck-Institut für Molekulare Pflanzenphysiologie, Potsdam-Golm, 14476, Germany

**Keywords:** temporal synchronization, rhythmicity, metabolism, leaf ecophysiology, domestication *de novo*

## Abstract

Cells, tissues, and organs are characterized by harbouring complex systems allowing communication between one another. Plant domestication was demonstrated to have structured the circadian rhythms, while also synchronising flowering and metabolism. Here, we demonstrate that the domesticated tomato (*Solanum lycopersicum*) manifests more synchronized rhythmicity across the whole plant. Consequently, the leaf development program is more coordinated in this species than in its wild relatives, wherein *S. lycopersicum* young leaves develop slowly in comparison to mature leaves. Young leaves from wild tomatoes display higher photosynthesis than mature leaves, while large metabolite accumulations occur across plant segments. Consequently, the diel metabolite levels are rather similar between young and mature leaves in the wild tomato *S. pennellii*, whereas the expression patterns for circadian clock genes are widely contrasting between both leaves. We further demonstrated that additions of genes related to domestication into the wild tomato *S. pimpinellifolium* appear to synchronize the development of young and mature leaves to be rather similar to that observed for *S. lycopersicum*. Collectively, the strengthening of inter-organs relationships on domesticated tomato indicates a synchronized biology, which is most likely fundamental to explaining its elevated yield.

## INTRODUCTION

Historically, human beings have been selecting plants to ensure their food security and support population growth, leading to drastic alterations in plant development. Most plant traits are affected by timing, demanding accurate mechanisms to record and respond to time. Accordingly, timekeeping mechanisms monitor geophysical time and record coincidences, which adjust the organism’s development and frequently promote the anticipation of future environmental conditions. A single organism may have diverse timekeepers working either independently or in synchronicity, for example, clocks associated with development, circadian rhythms, and metabolic oscillations (Zhu et al., 2020; Park, et al., 2012; Sanchez and Kay, 2016; Papagiannakis et al., 2017; Siqueira et al., 2018). As such, the developmental clock defines the precise timing to induce cell fates related to the growth of organs and subsequently shifting their morphology (Zhu et al., 2020). The circadian clock, in turn, represents the timekeeper that synchronises the endogenous system of the organism following “solar time”, to regulate oscillating rhythms according to a period of 24 hours (Sanchez and Kay, 2016). Furthermore, autonomous metabolic oscillations may occur across diverse growth conditions independently from other oscillatory systems, characterising the so-called metabolic clock (Papagiannakis et al., 2017; Siqueira et al., 2018). Remarkably, it remains unclear as to why biological clocks in modern crops are characterized by triggering the phenomena of flowering synchronization, circadian clock deceleration, and broader metabolic adjustments than their wild relatives.

Domestication is assumed to have mainly modified the developmental clock of wild plants, altering the shoot apical meristem (SAM) in a manner that shapes architecture and synchronizes the flowering time of plants (Doebley et al., 2006). The SAM maturation clock in domesticated tomatoes (*Solanum lycopersicum*) shows a more advanced maturation status than in tomato wild species, culminating in a fast transition from inflorescence meristem to floral meristem (Park et al., 2012). Notably, photoperiod is a dominant factor orientating flowering induction and the ability to sense changes based on day length (Wahl et al., 2013), which allowed the spread of most crop species worldwide. Varieties of cultivated tomatoes are insensitive to day length, while their wild relatives are not, and this difference is partially explained by variations in the *cis*-regulatory region of the gene *Self-Pruning 5G* (*SP5G*) (Soyk et al. 2017). Under long-day conditions, the induction of the *SP5G* gene arrests the flowering of wild tomatoes, whereas the repression of *SP5G* expression under long-day conditions enables the flowering of domesticated tomatoes (Soyk et al. 2017). Furthermore, photoperiod is a central factor governing the rhythmicity of circadian clocks, whereas tomato domestication selected plants with a slower circadian clock (Müller et al., 2016). The reproductive performance of *S. lycopersicum* is specifically improved over long days owing to this selection, allowing their cultivation under summer days at the higher latitudes of the northern hemisphere (Müller et al., 2016). A key component connecting developmental and circadian clocks appears to be metabolism, as revealed by metabolic signatures which are assumed to orientate the domestication of some species, including adzuki bean (*Vigna angularis*) and barley (*Hordeum vulgare*) (Yang et al., 2015; Pankin et al., 2018). Singularities in temporal biology are expected to occur across plant species, and as such temporal coincidences among developmental, circadian, and metabolic clocks must occur to enable the improved yield of domesticated tomatoes in comparison to their wild relatives.

Communication across tissues of complex biological systems occurs due to cell-to-cell relationships, contributing to multicellularity structuration in both animals and plants. Accordingly, mice (*Mus musculus*) submitted to a high-fat diet had losses in the temporal coincidences of circadian metabolism among eight distinct tissues, promoting aberrant cell proliferation and growth of these tissues (Dyar et al. 2018). Notably, mice liver exhibits an independent circadian clock oscillating autonomously from all other clocks, while this clock is dependent on the light/dark cycles to sustain the rhythmicity (Koronowski et al., 2019). In good agreement, each leaf tissue exhibits a particular clock that expresses differential contributions to the global leaf circadian clock, and the rhythmicity is asymmetrically coupled across tissues, ensuring correct leaf development in *Arabidopsis thaliana* (Endo et al., 2014). Recently, we suggested that domestication was an agent setting biological clocks on cultivated species, reducing the heterogeneity among distinct tissues (Siqueira et al., 2022). It seems reasonable to posit that this may have contributed to photoperiod adaptation and synchronized flowering in domesticated plants (Xiang et al., 2022; Siqueira et al., 2022). To reveal potential synchrony among biological clocks in wild and domesticated tomatoes, we combined physiological, metabolic, and genetic assays to demonstrate differences among these species. Our results revealed that domesticated tomato has synchronized circadian rhythms in both cotyledons, a fact that is not observed for the wild tomato *S. pimpinellifolium*. We additionally demonstrated that metabolism and gene expression are rather integrated between leaves of domesticated tomatoes, which appears to be a factor to help us in explaining their elevated yield in comparison with their wild relatives.

## MATERIALS AND METHODS

### Plant material and growth conditions

Tomato seeds were provided by Professor Dr. Lázaro E. P. Peres (Escola Superior de Agricultura Luiz de Queiroz da Universidade de São Paulo (ESALQ/USP), Brazil). Seeds from *Solanum lycopersicum* (Cv. M82), *S. pimpinellifolium* (LA LA1589), *S. habrochaites* (LA1777), *S. neorickii* (LA2133), and *S. pennellii* (LA716). We additionally selected introgression lines (IL5-4, IL9-2-6, and IL9-3) harbouring genome fragments of *S. pennellii* in the genetic background M82 (*S. lycopersicum*), which were obtained from Zamir *et al*. (1995). In addition, we performed assays with *S. pimpinellifolium* lines harbouring editions in six loci related to tomato domestication, as described by Zsögön *et al*. (2018). Seeds were surface-sterilized with 5% sodium hypochlorite for 10 min and then washed with running distilled water and subsequently sowed in a tray with a commercial substrate (Tropstrato HT^©^). Two weeks after the sowing, seedlings were transferred to 400 mL pots (one seedling per pot) containing the same substrate used for seedling production but supplemented with 1 g of NPK 10–10–10 and 4 g dolomitic limestone (Ca + Mg) per L of the substrate (Pino-Nunes et al. 2018). Plants were grown in a controlled greenhouse located in Viçosa (20°45′S, 42°50′W, 650 m above sea level), Brazil, with a mean temperature of 28 °C, 12.0 hours (winter/spring/summer) of photoperiod, and a minimum of 600 μmol photons m^−2^ s^−1^.

Plants were watered regularly and throughout the entire growth period were maintained under naturally fluctuating conditions of light intensity, temperature, and relative air humidity. All physiological, and biochemical parameters analysed in the experiments were performed on the third and fourth leaves when source leaves were completely expanded, which occurred for 4-week-old plants. Additionally, each experiment was repeated at least three times (even in different growth facilities) with similar phenotypes observed each time. Throughout the experiment, the plants were grown under naturally fluctuating conditions of temperature and air relative humidity and were fertilized, as necessary. The pots were randomized periodically to minimize any variation within each light environment. For biochemical and molecular analyses, leaf samples were collected at different times during the day, before flash freezing in liquid nitrogen (N_2_) and subsequent storage at −80 °C, where it remained until analysis. The means presented in the tables and figures were obtained from five independent replicates per treatment of single plant experimental plots (one plant per pot). The experiments were repeated at least three times with similar phenotypes observed each time.

### Cotyledon movements and circadian rhythm determination

Seedlings were grown in controlled conditions of a growth chamber for four days under cool white fluorescent tubes (~100 μmol m^−2^ s^−1^) under 12h light/ 12h dark and 20:18 °C temperature, which was transferred subsequently to constant light and temperature (25 °C). A polystyrene ball was attached to the tip of the cotyledons of each seedling by using petroleum jelly according to Müller *et al*. (2016).

### Growth analyses

Growth parameters were determined in 4-week-old plants by measuring leaf area, length, and mass as well as the specific leaf area (SLA). Leaf area was measured using a scanner (Hewlett Packard Scanjet G2410, Palo Alto, California, USA) and processing the resulting images on ImageJ. SLA was measured as described previously (Hunt, 2002). At the end of the experiment, plants were harvested by cutting the segments above ground level in the sense of the third leaf for the fourth leaf, thus separating leaflets, petioles, and stem.

### Measurements of gas exchange and chlorophyll fluorescence

Gas exchange parameters were determined simultaneously with chlorophyll a (Chl a) fluorescence measurements by using an open-flow infrared gas exchange analyser system (LI-6400XT; LI-COR Inc., Lincoln, NE) equipped with an integrated fluorescence chamber (LI-6400-40; LI-COR Inc.). Instantaneous gas exchanges were measured after 1-hr illumination during the light period under 1,000 μmol m^−2^ s ^−1^ at the leaf level (light saturation) of photosynthetically active photon flux density (PPFD). The reference CO_2_ concentration was set at 400-μmol CO_2_ mol^−1^ air. All measurements were performed using the 2-cm2 leaf chamber at 25°C, as well as a 0.5 stomatal ratio (amphistomatic leaves), and leaf-to-air vapour pressure deficit was kept at 1.2 kPa, and the amount of blue light was set to 10% PPFD to optimize stomatal aperture. Briefly, the initial fluorescence emission (F0) was by illuminating dark-adapted leaves (1 h) with weak modulated measuring beams (0.03 μmol m^−2^ s^−1^). A saturating white light pulse (8,000 μmol m^−2^ s^−1^) was applied for 0.8 s to obtain the maximum fluorescence. In light-adapted leaves, the steady-state fluorescence yield was measured with the application of a saturating white light pulse (8,000 μmol m^−2^ s^−1^) to achieve the light-adapted maximum fluorescence (Fm′). According to Genty et al. (1989), ΦPSII represents the number of electrons transferred per photon absorbed in the PSII, and the electron transport rate (J_flu_) was calculated. Dark respiration (*R_d_*) was measured after 2 hours in the dark period (at night), using the same gas exchange system described above, and it was divided by two (Rd/2) to estimate the mitochondrial respiration rate in the light (RL). Determination of mesophyll conductance (*g_m_*), maximum rate of carboxylation (*V_cmax_*), maximum rate of carboxylation limited by electron transport (*J_max_*), and photosynthetic limitations The CO_2_ concentration in the carboxylation sites (*C_c_*) was calculated according to (Harley et al., 1992). Briefly, this method uses the values of *A_N_*, *g_s_*, *g_m_*, *V_cmax_* and *C_c_*, and permits the partitioning into the functional components of photosynthetic constraints related to stomatal (L_s_), mesophyll (L_m_), and biochemical (L_b_) limitations.

### Determination of metabolite levels

Tomato segments were harvested and immediately frozen in liquid N2 and stored at −80°C until further analysis. For leaves, all sampling procedures were carried out on the same leaf used for the analysis of gas exchange and fluorescence. Leaf samples were harvested at different time points along the light-to-dark cycle (6:00, 12:00; 18:00, 00:00, and 6:00). The extraction of metabolite extraction was executed by rapid grinding in liquid N2 and immediate addition of the specific extraction buffer. Chlorophyll levels were determined according to Wellburn (1994), whereas glucose, fructose, sucrose, and starch were determined as described previously (Fernie et al., 2001). Total amino acids and soluble proteins were determined as previously described by Yemm *et al*. (1955) and Bradford (1976), respectively.

Other metabolites were quantified according to gas chromatography associated with the mass spectrometry (GCMS) protocol (Roessner et al. 2001; Lisec et al. 2006). We used approximately 50 mg fresh weight to perform the extraction procedure, by using 1 mL of methanol and shaking (800 rpm) at 70 °C for 15 min; being that 60 μL of ribitol (0.2 mg mL^−1^) was added as an internal standard. Subsequently, derivatization and sample injection were performed as previously described by Lisec *et al*. (2006). Chromatograms and generated mass spectra were evaluated using the software TagFinder (Luedemann et al. 2011), using a reference library from the Golm Metabolome Database (Kopka et al., 2004), and the identification and annotation of the detected peaks followed the recommendations for reporting metabolite data described in Fernie *et al*. (2011).

### Gene expression

Total RNA was isolated from 50 mg of leaves using the TRIzol reagent (Invitrogen®), according to the manufacturer’s instructions. The integrity and amount of RNA were checked on 1% (w/v) agarose gel, and the concentration was measured before and after DNase I digestion (RQ1 RNase free DNase I, Promega®) using spectrophotometry (OD_260_). DNase-treated RNA (2 μg) was used for cDNA synthesis using the SuperscriptTM III reverse transcriptase (Invitro-gen®, Darmstadt, Germany) according to the manufacturer’s recommendations. Gene expression was accessed using real-time PCR (qRT-PCR) (Step One PlusTMReal Time PCR System, Applied Biosystems, CA, USA) with the SYBR green fluorescence detection (Applied Biosystems®), using the Platinum® SYBR® Green qPCR SuperMix-UDG with ROX kit. Data analyses were performed exactly as described by Caldana et al. (2007). qRT-PCR was performed according to the following temperature alternations: denaturation of double-stranded DNA and enzyme activation (95 °C, 30 s, 1x); denaturation of double-stranded DNA (95 °C, 3 s, 40x), and finally, primer’s annealing and extension of fragment by polymerase enzyme (58 °C, 30 s, 40x). Melt curve reaction: 95 °C, 15 s, 60 °C for 1 min, and 95 °C for 15 s (1x). The average CT of the ACTIN gene (reference control) was used for the relative expression analyses. The analyses were performed in duplicate in each PCR run, using 4 replicates for each genotype, and their mean cycle threshold was used for relative normalized expression analyses.

### Statistical analyses

The experiments were randomized and designed with a minimum of three biological replicates of each treatment. Furthermore, experiments to describe phenotypes were repeated at least three times. Data were statistically tested for normality and subsequently examined using ANOVA (*P* < 0.05). Differences in the means (*P* < 0.05) displayed in figures and tables were examined by Student’s *t-test*. All statistical analyses were performed using R statistical software (www.rproject.org).

## RESULTS

### The similar circadian rhythms at cotyledons of Solanum lycopersicum

Tomato domestication decelerated the circadian rhythms of cultivated tomato (Müller et al., 2016; Müller et al., 2018), whereas these rhythms could be variable across tissues of wild plants. Each circadian clock seems to contribute to specific processes in the leaf, and differently from the centralized circadian clock of mammals, the plant circadian clock seems not exhibit a clear centralization (Shimiziu et al., 2015). Nevertheless, shoot apex clocks seem to have the capacity to synchronize circadian rhythms across plant structures, affecting rather distant organs such as circadian oscillations of roots (Takahashi et al., 2015; Chen et al., 2020). Despite certain imprecision regarding the centralization of circadian rhythms in plants, it remains the consensus that circadian misalignment is greatly deleterious for the fitness of both mammals and plants (Koronowski and Sassone-Corsi, 2021; Sorkin and Nusinow, 2021). Based on this consensus, we decided to assess circadian rhythms in domesticated and wild tomatoes.

The exposure of the two *A. thaliana* cotyledons to constant light revealed contrasting circadian rhythms for both over several days, indicating the independence of each cotyledon clock (Thain et al., 2000). Therefore, we assessed circadian rhythms in cotyledon pairs of *S. lycopersicum* and *S. pimpinellifolium*, which revealed contrasting rhythms only for the wild species (Fig. 1A-B). Accordingly, the difference for the circadian period was 0.8 h between the cotyledons of *S. pimpinellifolium*. Thereafter, we turned our attention to identifying the developmental and physiological consequences of the absence of differences in circadian rhythms in the domesticated species.

**Figure 1.**
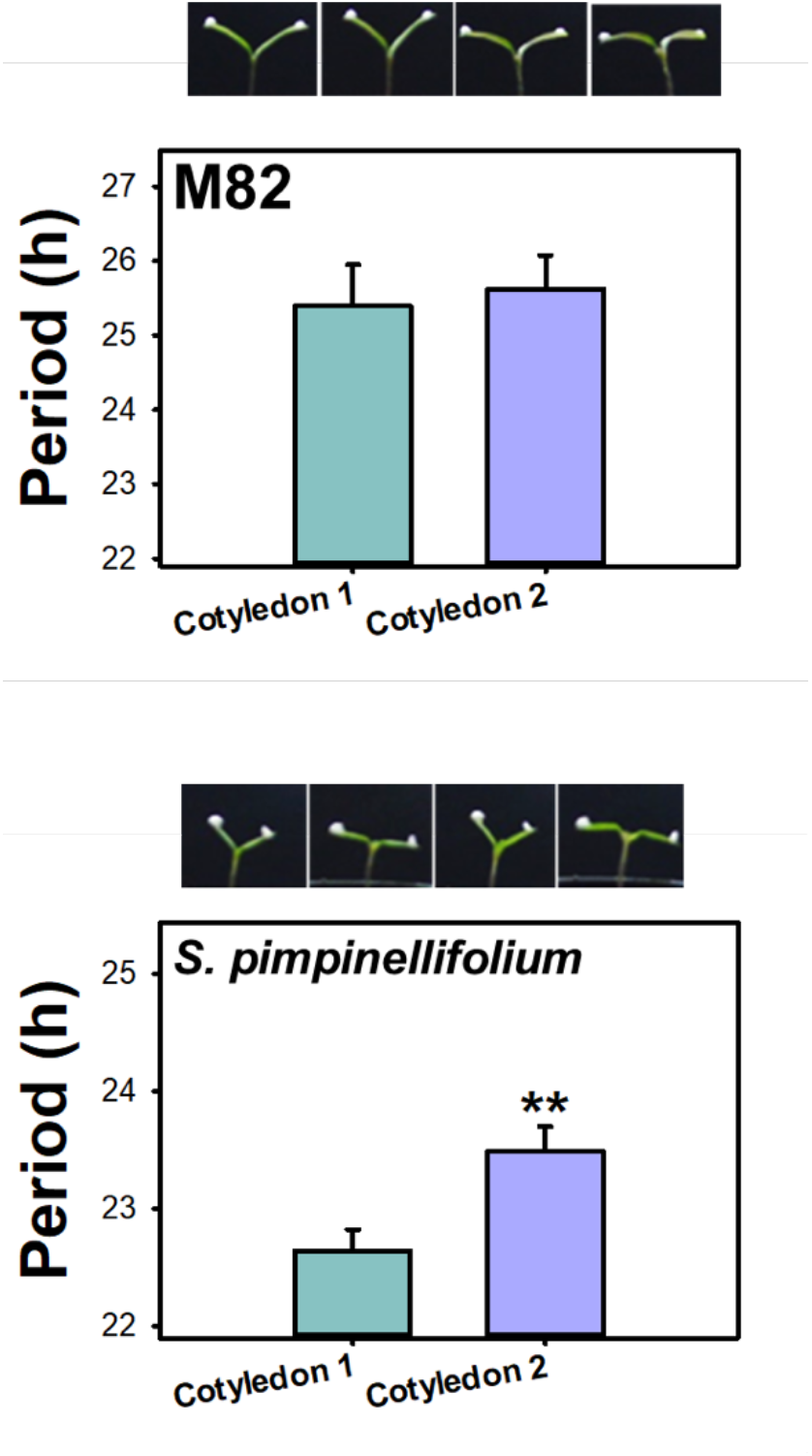
The circadian rhythms of cultivated tomatoes operate synchronously. Images showed cotyledons positions at 12, 36, 60 and 84 hours upon exposure to constant light conditions. Circadian period of the cotyledons from *Solanum lycopersicum* (cv. M82) and *Solanum pimpinellifolium*. Data presented are mean ± SE (n = 8), and an asterisk (*) indicates different values that were determined by the two-sided Student’s *t*-test to be different (*P* < 0.05) between cotyledon 1 and cotyledon 2.

### The developmental hierarchy of leaves in domesticated tomato

Most of the domestication and breeding of plants were performed based on reproductive yield, given that humans most likely selected plants according to desirable fruits and grains. Even though leaf shape was recently demonstrated to be an excellent predictor for fruit production and quality in domesticated tomatoes (Rowland et al., 2020), the role of leaves to ensure elevated crop productivities was rather neglected. In agreement with this observation, latitude and its relationship with day-length are closed associated with tomato domestication, allowing tomato production under long days in the summer of the Northern hemisphere (Müller et al., 2016; Soyk et al., 2017; Xiang et al., 2022), which may have differently affected tomato organs. We thus decided to investigate the major tomato traits affected by latitude using the data from the germplasm bank of Universidade Federal de Viçosa, Brazil (www.bgh.ufv.br). For this purpose, we mapped the relationships among latitude with growth traits of distinct organs in *S. lycopersicum* genotypes. Our *in silico* assays revealed that leaf traits are correlated with latitude in approximately 100 tomato genotypes from diverse regions of the Earth (Supplementary Fig. S1-2).

We next decided to turn our attention to leaf development and investigate whether *S. lycopersicum* could have a synchronized development of leaves. Although canopies of cultivated plants have homogenous leaves, they capture light heterogeneously (Long et al., 2015). Indeed, analyses of the leaf developmental trajectories in *S. lycopersicum* revealed an uneven photosynthetic competence, which spreads heterogeneously across leaf zones (Martinez et al., 2021). Investigating the leaf development in wild tomatoes, it a complex behaviour in leaf series was revealed, in which shade avoidance is ephemeral and leaf length and area are widely variable (Chitwood et al. 2012a). These traits tend to negatively correlate with the shade avoidance index during early development but positively correlate at later stages of leaf development (Chitwood et al. 2012a). Thus, wild tomatoes exhibit unequal shade avoidance along a leaf developmental series, whereby each leaf might exhibit a distinct degree of shade avoidance (Chitwood et al. 2012a; Chitwood et al. 2012b). Domesticated and wild tomatoes are characterized by an ephemeral shade avoidance that is not continuous over leaves (Chitwood et al. 2015). Thus, complex developmental patterns were visualised in leaves of a single plant, which due to a differential light exposure of them most likely promotes differential shade avoidance across plant regions (Chitwood et al. 2015). By analysing canopy and leaf patterns, we observed that the developmental plasticity seems to be higher in wild tomatoes in comparison with domesticated ones (Supplementary Fig. S3), indicating a leaf shape homogenization associated with tomato domestication. Thus, we postulated that a potential developmental overlap may exist covering the third and fourth leaves of wild (*S. pimpinellifolium*, *S. habrochaites*, *S. neorickii*, and *S. pennellii*) tomatoes which does not occur in the domesticated species. To reduce the impacts of variations in leaf shape and length, we performed analyses mainly on older leaflets from third (L3) and fourth (L4) leaves, which allowed to us assess both developmental and physiological competence from 15 to 35 days after sowing (DAS). Intriguingly, the same developmental pattern was observed for all wild tomatoes, wherein the L3 had a higher leaf area than L4 only at the first observations, in general around 15 or 20 DAS (Figure 2A). In sharp contrast, the domesticated tomato was always characterized as L3 displaying a larger leaf area than L4 across all experimental observations (Figure 2A), suggesting the occurrence of strict developmental coordination in *S. lycopersicum*.

**Figure 2.**
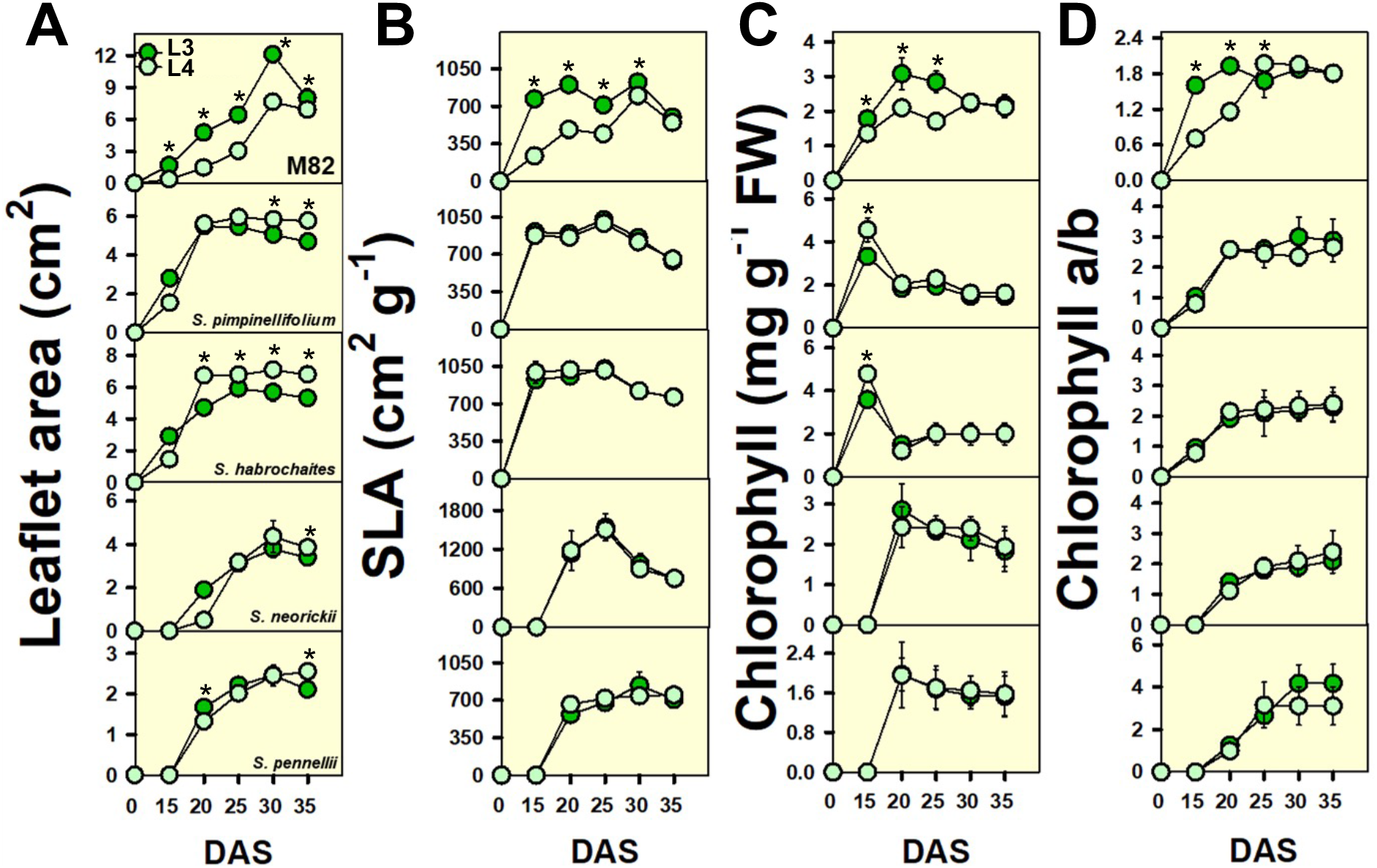
Developmental and physiological traits over development from domesticated and wild species of tomatoes. Leaflet traits were assessed in older leaflets of third (L3, dark-green) and fourth (L4, pale-green) leaves from domesticated tomato (*Solanum lycopersicum* cv. M82), *S. pimpinellifolium*, *S. habrochaites*, *S. neorickii*, and *S. pennellii*. **A**, Leaflet area (cm^2^); **B**, SLA: specific leaf area (cm^2^ g^−1^); **C**, Chlorophyll (mg g^−1^ fresh weight); **D**, ratio chlorophyll a/b. Data presented are mean ± SE (n = 7), and an asterisk (*) indicates different values that were determined by the two-sided Student’s t-test to be different (*P* < 0.05) between L3 and L4 leaflets in the specific point. DAS: days after sowing.

Specific leaf area (SLA) is an important physiological trait describing the plant’s ability to intercept photosynthetic irradiation. Commonly, older leaves exhibit a lower SLA than young leaves, denoting a better photosynthetic ability and a more advanced development due to cell differentiation *status* (Baldazzi et al., 2013). SLA was essentially invariant between L3 and L4 over the early development of wild tomatoes, whereas it was different between L3 and L4 during the 15 to 30 DAS in *S. lycopersicum* (Figure 2B). The physiological maturity of a leaf occurs when it reaches the status of the source of photoassimilates for other plant organs. Accordingly, chlorophyll content is an excellent indicator of physiological maturity revealing when a leaf is no longer a sink but rather a source of sugars. We thus measured chlorophyll levels over early development, finding a similar pattern to that observed for leaflet area and SLA, with differences practically absent between L3 and L4 even in the early developmental stages of wild tomatoes (Figure 2C). Accordingly, only domesticated species displayed differences in chlorophyll levels and chlorophyll *a*/*b* ratio (Figure 2C-D). Decreases in chlorophyll levels exert a direct role in the reduction of the number of lateral branches, which seems to modulate shoot branching and alter energy balance (Khangura et al., 2020). Likewise, we next determined the levels of sugars related to energy homeostasis, wherein it was observed only for *S. lycopersicum* highest sugar levels present in L3 than in L4 during the early stages of the leaf development (Supplementary Fig. S4). Over development the glucose and fructose levels were increasing in L4, wherein higher levels of these compounds were found in later developmental stages for L4 in comparison to L3, a fact that was not observed for sucrose (Supplementary Fig. S4). Meanwhile, we did not observe a clear pattern for sugar levels in wild tomatoes, and as such L3 and L4 frequently had similar levels of glucose, fructose, and sucrose regardless of the developmental stage. Altogether, these findings suggest that development and metabolism are likely coordinated and synchronized in *S. lycopersicum* while wild tomatoes exhibited clear independence between L3 and L4.

We further performed light and CO_2_ curves using *S. lycopersicum* cv. M82, *S. pimpinellifolium*, and *S. pennellii* to compare gas exchange responsiveness between L3 and L4. Net photosynthesis (*A_N_*) is homogenous in leaflets of M82 and *S. pimpinellifolium*, whereas the L4 of *S. pennellii* had higher *A_N_* than L3 in most levels of light and CO_2_ (Figure 3). Attempting to understand the differences observed, we split seven segments from L3 to L4 of M82 and *S. pennellii* to map metabolic alterations across these segments. We observed that M82 was metabolically invariant across segments, while *S. pennellii* exhibited remarkable differences in metabolite accumulation specifically in metabolites related to sink function including sucrose, citrate, and GABA (Figure 3). Collectively, these results indicated that wild tomato leaflets reach faster maturity becoming independent organs which means that they can be self-sufficient at early development, which contrasts with the observations for the domesticated tomato.

**Figure 3.**
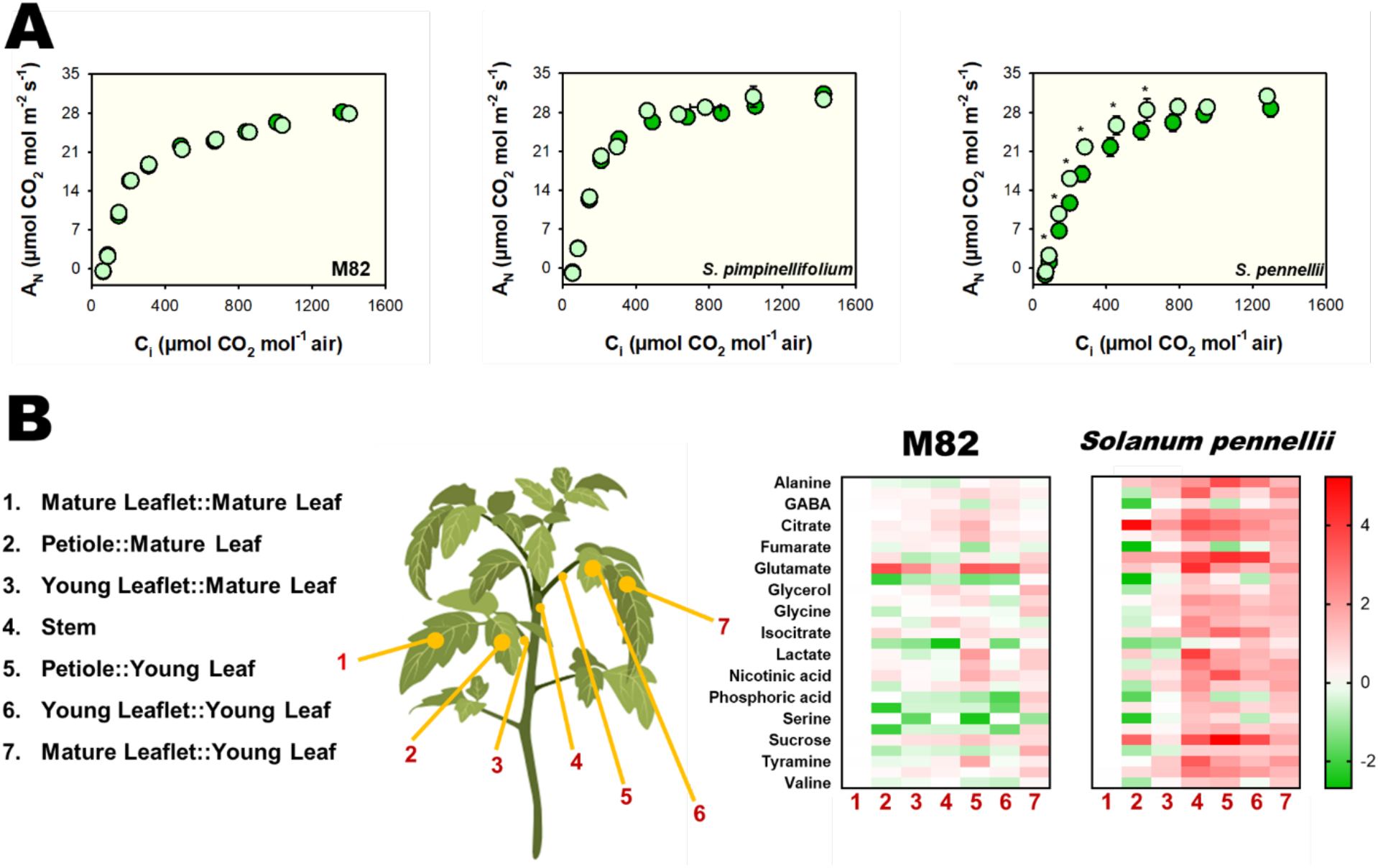
Net photosynthesis (*A*_N_) curves in response to sub-stomatal (Ci) CO_2_ concentration in domesticated and wild tomatoes. **A**, *A*_N_/*C*_i_ curves of *Solanum lycopersicum* (Cv. M82), *S. pimpinellifolium*, and *S. pennellii* determined in leaflets of (dark-green) and fourth (pale-green) leaves. Data are means ± SE (n = 7), and an asterisk (*) indicates different values that were determined by the two-sided Student’s t-test to be different (*P* < 0.05) between the third and fourth leaflets. **B**, Metabolite levels at the middle day were assessed in seven segments. Samples were taken from the third and fourth leaves from the apex of 4-week-old plants. Data are means ± SE (n = 5).

Physiological traits seemingly play a more important role in determining high rates of leaf growth than morphological features, with domesticated tomatoes being more impacted by the variance in these traits than wild tomatoes (Conesa et al., 2017). Thus, we investigated leaflet physiology in M82, *S. pennellii*, and introgression lines (ILs) harbouring *S. pennellii* genomic fragments. These ILs allowed the identification of several quantitative trait loci (QTLs) (Eshed and Zamir, 1995), including the identification of regions intimately associated with tomato domestication, branching, flowering, photoperiodism, and circadian clocks. In this sense, IL5-4 has the genomic fragment harbouring the *SP5G* gene that regulates architecture and flowering according to photoperiodic responses whereas IL9-2-6 and IL9-3 harbours the genes that promote the circadian clock deceleration (Müller et al., 2016; Soyk et al., 2017). By investigating the L3 and L4 development of *S. lycopersicum* (cv. M82), IL5-4, IL9-2-6, IL9-3, and *S. pennellii*, we observed a similar developmental window for these leaflets in ILs and *S. pennellii* (Supplementary Table 1). Thus, around 35 DAS, the area and mass of leaflets were similar between L3 and L4 of the same plant in the ILs and in wild species, yet this relationship does not occur in *S. lycopersicum* in which L3 displays a higher area, SLA and mass than L4 (Supplementary Table 1). By contrast, ILs and wild tomatoes showed similar SLA for leaflets L3 and L4, describing a similar ability to intercept radiation (Supplementary Table 1), which suggests a similar developmental trajectory. In consonance with these developmental coincidences, we next decided to investigate whether ILs and *S. pennellii* would exhibit contrasting physiologies for each leaflet.

### Heterogeneous physiology but similar metabolism between leaves of S. pennellii

Tomato domestication resulted in remarkable changes in photosynthesis-related genes, wherein an inverse correlation between transcripts related to leaf development and photosynthesis can be noted (Ranjan et al., 2016). Intriguingly, despite *S. pennellii* leaflets exhibiting the same SLA, a higher *A_N_* for L4 in comparison with L3 was observed (Supplementary Figure S6). Meanwhile, our results revealed that *S. pennellii* exhibited a differential *V*_*c*max_ for different leaves of the same plant, with the highest values for L4 than L3 (Supplementary Figure S6). In agreement with these data, a similar pattern for ILs and *S. pennellii* was observed for *A*_N_, *R*_d_, *P*_R_, *g*_m_, *Vc*_max_, and biochemical limitations, supporting the notion of weak source-to-sink relationships in these plants (Supplementary Fig. S6-7). Taken together, our findings suggested that these genomic regions might regulate the metabolic synchronization between L3 and L4. To further elucidate this, we decided to perform metabolite analyses on these leaflets in an attempt to identify the source-to-sink patterns.

Despite the higher *A_N_* at L4, the diel metabolism of glucose, fructose and sucrose revealed few differences between L3 and L4 for ILs and *S. pennellii* (Figure 4 and Supplementary Figure S7). Interestingly, the coupling between organism size and metabolic rates displays few relationships with the selection of plants over domestication (Milla et al., 2018), which may be due to the weak inter-tissue metabolic relationships. To summarize, the atemporal coordination of sucrose/starch metabolism between L3 and L4 on ILs and *S. pennellii* seems to describe a rather weak inter-leaf relationship. Metabolite diversity in wild and domesticated *Phaseolus vulgaris* revealed that tissue specificity is the major factor affecting the metabolite dataset, yet wild genotypes display more metabolite diversity (Souza et al., 2019). Meanwhile, considering a single tomato plant, metabolite specialization across different tissues seems to result from differential epigenetic regulation that in turn promotes differential gene expression in mature and young organs (Zhong et al., 2013). Taking the findings of these two studies together, it seems reasonable to anticipate that gene expression might explain the contrasting metabolic regulation at L3 and L4 of M82 and *S. Pennellii*.

**Figure 4.**
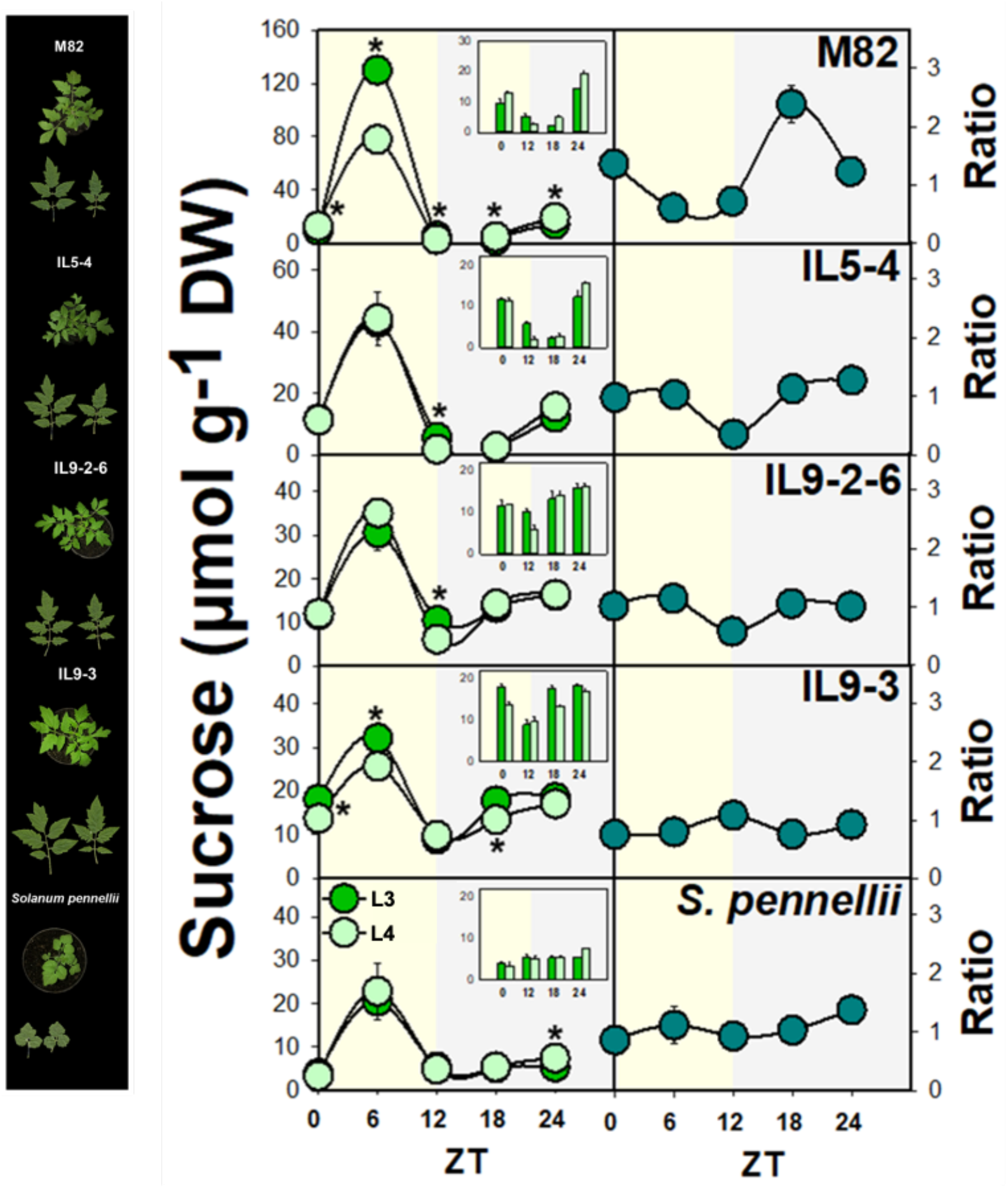
The diel regulation of sucrose metabolism on tomatoes. Older leaflets of (L3, dark-green) and fourth (L4, pale-green) leaves from domesticated tomatoes (Cv. M82), introgression lines (IL) and wild tomato *Solanum pennellii* were analyzed, as represented at left, while the right shows sucrose levels and the ratio between the fourth and third leaflets. Sucrose levels were assessed in leaflets harvested every 6 hours over a diurnal cycle. Data are means ± SE (n = 5), and an asterisk (*) indicates different values that were determined by the two-sided Student’s t-test to be different (*P* < 0.05) between L3 and L4 leaflets in the specific point. Yellow bars indicate the light period while grey bars indicate the dark period. DW: Dry weight.

### Transcriptional regulation of leaves

Transcriptional variation has been widely used to monitor circadian rhythms over the past years. For this reason, we next assessed the gene expression of M82 and *S. pennellii* growing under natural light/dark cycles to identify potential synchronicity between L3 and L4. *TIMING OF CAB EXPRESSION1* (*TOC1*) is a pivotal component in circadian clock regulation since it represses and is repressed by *LATE ELONGATED HYPOCOTYL* (*LHY*) (Nohales and Kay, 2016). Under natural conditions *TOC1* expression at shoots is sufficient to ensure plant survival under drought while its expression only in roots did not promote a similar effect (Valim et al., 2019), supporting the notion of clock communication across plant organs. We observed contrasting expressions for *TOC1* during all diel period on L3 and L4 of M82 and *S. pennellii* (Figure 5). Circadian period decreases with leaf age progression, and the shorter period of older leaves is linked with *TOC1* expression in *A. thaliana* (Kim et al., 2016). Thus, the expression of *TOC1* on *S. pennellii* was always higher in L3 than L4, whereas it was higher in L4 than L3 under darkness in M82, with an almost similar pattern observed for *LHY* expression (Figure 5), suggesting an expression hierarchy based on the development for the wild species. *GIGANTEA* (*GI*) is a positive regulator of TOC1 and an integrator between the circadian clock and development that contributes to floral induction (Nohales and Kay, 2016). We found a homogenous *GI* expression between L3 and L4 on M82, whereas larger expressions at L3 than at L4 were found in *S. pennellii* (Figure 5). Taken together, the expression of *TOC1*, *LHY*, and *GI* seem to indicate a coordinated regulation of the circadian clock at the leaflet level in the domesticated species, while in wild relatives a more particular transcriptional regulation for clocks of each leaflet was observed.

**Figure 5.**
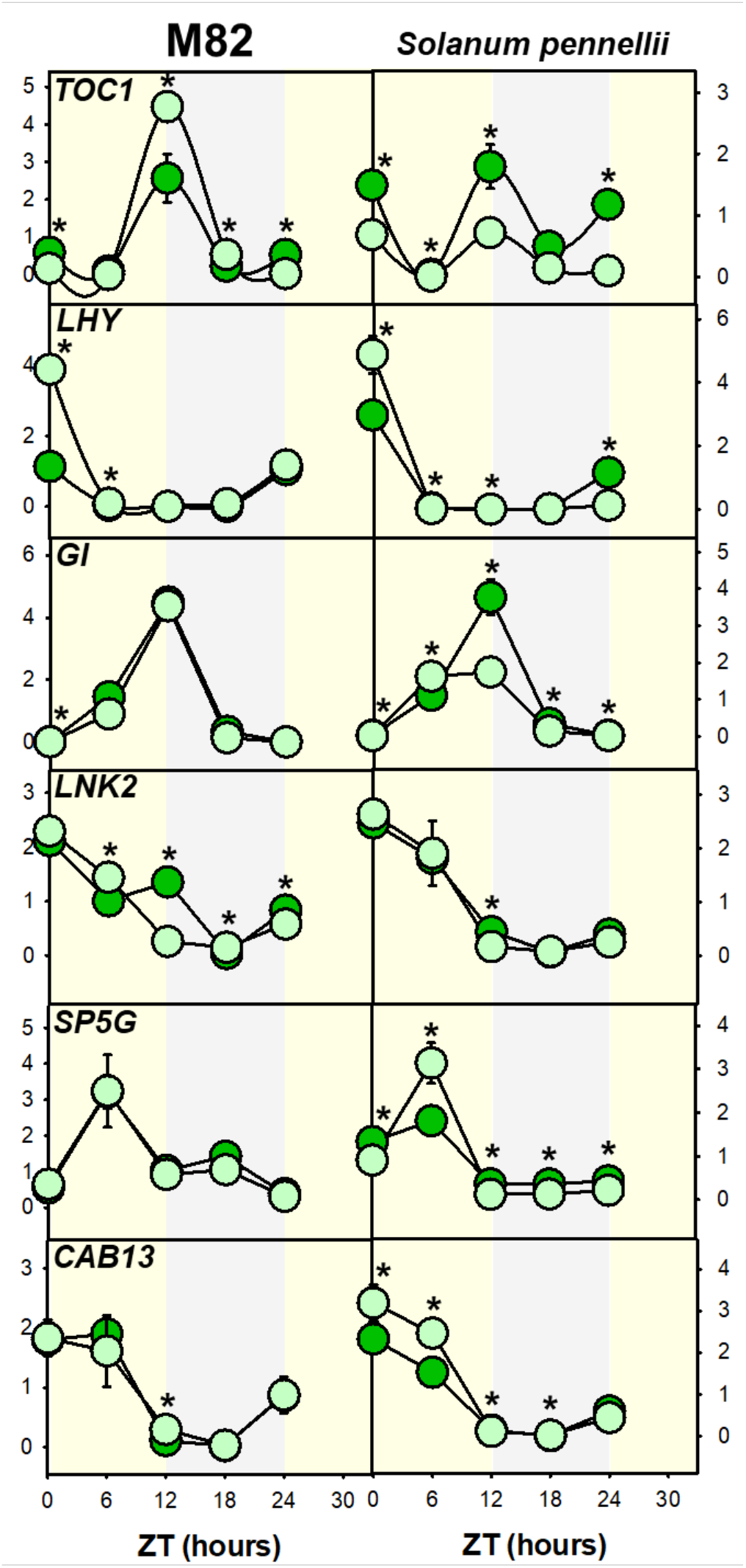
Expression profile of genes associated with tomato domestication in *Solanum lycopersicum* cv. M82 and *S. pennellii*. Diurnal oscillations of transcript levels were determined in leaflets of the third (dark-green) and fourth (pale-green) leaves. An asterisk (*) indicates different values that were determined by the two-sided Student’s t-test to be different (P < 0.05) between the third and fourth leaflets. Data represent the average expression of three biological replicates ± SE. TIMING OF CAB EXPRESSION 1 (TOC1); LATE ELONGATED HYPOCOTYL (LHY); GIGANTEA (GI); NIGHT LIGHT-INDUCIBLE AND CLOCK-REGULATED 2 (LNK2); SELF PRUNING 5G (SP5G); CHLOROPHYLL AB BINDING PROTEIN 13 (CAB13).

Circadian clock deceleration in domesticated tomatoes occurred due to selective pressures on genes EID1 and *NIGHT LIGHT-INDUCIBLE AND CLOCK-REGULATED 2* (*LNK2*), with both genes functioning in light signalling and flowering (Müller et al., 2016; Müller et al., 2018; Rugnone et al., 2013). A sharp contrast was observed for *LNK2* expression in domesticated tomato, wherein L3 and L4 expression were distinct over the diel period but the expression was virtually invariant in *S. pennellii* (Figure 5). In plants, *SP5G* programs seasonal changes in accordance with day length, exhibiting a diel expression that generally culminates in flowering repression in wild tomatoes under long days (Soyk et al., 2017). *SP5G* expression was similar in both leaflets of M82, and widely contrasting in L3 and L4 of *S. pennellii* across the diel period (Figure 5), supporting, at least in part, the notion of a particular developmental programming in each leaflet. *Light Harvesting Chlorophyll a/b Binding protein 13* (*CAB13*) obeys circadian dynamics, playing roles at photoinhibition mitigation and tolerance to constant light in wild tomatoes (Velez-Ramirez et al., 2014). No differences for *CAB13* between L3 and L4 from M82, whereas in *S. pennellii* L4 usually displayed higher *CAB13* expression when compared with L3 (**Figure 5**), supporting the notion of a differential photosynthetic regulation between L3 and L4 of wild species.

#### De novo Domestication, a promising strategy to re-structure synchronization of tomatoes

We next explored the de novo domestication *de novo* concept, performing assays with *S. pimpinellifolium* multiplex lines harbouring additions of multiple different genes related to tomato domestication. Complex genetic features are associated with the tomato domestication syndrome linking, for example, the fruit flavour with productivity (Alseekh et al., 2021). In addition, the *S. pimpinellifolium* genome is slightly more complex than that of *S. lycopersicum*, and therefore gene additions into domesticated tomatoes may result in unpredictable phenotypes (Alonge et al., 2020; Rodríguez-Leal et al., 2017). Since wild species endure hostile environments, their biological clocks integrate a plethora of abiotic stresses responses, and thus *de novo* domestication describes the introduction of domestication genes and/or re-domestication into wild species, representing a promising strategy to develop crop ideotypes (Markham and Greenham, 2021; Zsögön et al., 2018; Fernie and Yan, 2019; Yu et al., 2021). To exemplify the potential of this new concept, increased fruit size and number, as well as a higher nutritional value, were observed in engineered *S. pimpinellifolium* lines (Zsögön et al., 2018). By growing some of these engineered lines side-by-side, we verified that the gene additions also resulted in the rewiring of developmental coincidences between L3 and L4 (Figure 6). Briefly, the edited lines displayed the shortest leaflets for the young leaves, with a considerably lower area and biomass than the mature leaflets, as well as variations in SLA resembling source-to-sink relationships similarly to those previously observed for domesticated tomatoes.

**Figure 6.**
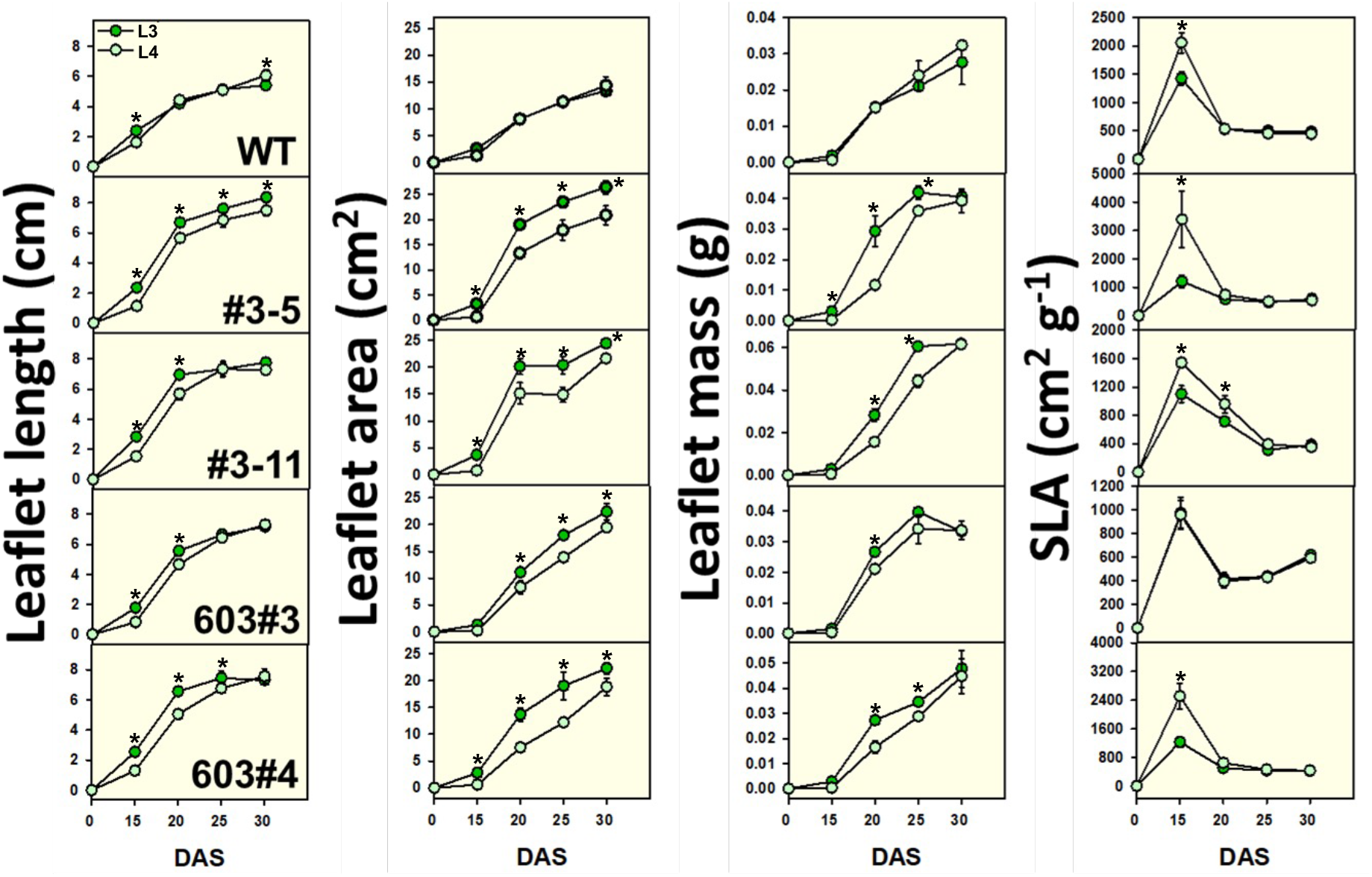
The establishment of leaf development hierarchy with domestication *de novo*. Leaflet traits were assessed in older leaflets of (L3, dark-green) and fourth (L4, pale-green) leaves from *Solanum pimpinellifolium* genotypes, wild-type (WT) and multiplex lines harbouring gene editions in key genes associated with domestication. Data presented are mean ± SE (n = 7), and an asterisk (*) indicates different values that were determined by the two-sided Student’s t-test to be different (*P* < 0.05) between L3 and L4 leaflets in the specific point. Leaflet length (cm); Leaflet area (cm^2^); Leaflet mass (g); SLA: specific leaf area (cm^2^ g^−1^); DAS: days after sowing.

## Discussion

Our study sheds light on a remarkable temporal difference in physiological and developmental regulation in leaves of wild and domesticated tomatoes, that ultimately indicates that the synchronization of the leaf biology can boost crop yield. Our findings are in consonance with observations in mammals that identified the independence of liver circadian rhythms from the central clock as well as the fact that the liver clock oscillates due to light/dark cycles (Koronowski et al., 2019). Thus, it was revealed that proper circadian clock communication elevated organism health, whereas highly caloric diets disrupt the coherence of circadian metabolism among tissues contributing to disordered growth in humans (Koronowski and Sassone-Corsi, 2021; Dyar et al., 2018). The situation in plants is rather complex since plants are largely dependent on the environment to orientate circadian rhythms, suggesting a different way in terms of inter-organ synchronization, since leaves may maintain their growth with less dependence on other organs/tissues. In this case, the exposure to constant light conditions promotes the desynchronization of circadian clocks whereas the increased multicellularity level precedes larger delays in anticipating outer cycles (Masuda et al., 2017; Muranaka and Oyama, 2016). Domestication of distinct plant species differentially shaped the rates of gas exchange. For example, whilst *Brassica rapa* has three morphotypes (leafy, turnip, and oilseed), domestication impacted *A*_N_ and stomatal conductance only in oilseed varieties (Yarkhunova et al., 2016). Furthermore, oilseed has a shorter circadian period coupled with a higher *Vc*_max_ than leafy and turnip morphotypes (Yarkhunova et al., 2016), whereas it was not investigated whether a temporal regulation across leaves of oilseed may exist. Domesticated tomato in turn display photosynthetic differences according to leaf age, wherein leaves with elevated age had much lower *A_N_* capacity and *R*_d_ than younger leaves (Xu et al., 1997). Accordingly, Rubisco activity (per g protein) is invariant across young and mature leaves of *S. lycopersicum*, whereas Rubisco total activity expressed per unit leaf area was lower in older than younger leaves (Xu et al., 1997), which seems to contribute to creating a photosynthetic hierarchy across leaves.

Autonomy to regulate genetic and metabolic traits is fundamental for organ development and thus synchronize tissues across organism with environmental variations. The misalignment of circadian relationships among tissues causes a desynchronized development prejudicing organism health (Dyar et al., 2018). Accordingly, each *A. thaliana* leaf tissue has particular circadian rhythms, and mesophyll and vascular clocks are the major drivers of the global leaf clock (Endo et al., 2014). This asymmetry among clocks may be synchronized and shoot apex clocks couples circadian rhythms of other organs, sending proteins to roots that increase or decrease the circadian period according to with temperature (Takahashi et al., 2015; Chen et al., 2020). The synchronization level among cells decreases with time progression and cell-to-cell communication probably enhances desynchronization, while individual plant cells under constant light manifest autonomous circadian oscillations (Masuda et al., 2017; Muranaka and Oyama, 2016). Circadian rhythms heterogeneity in intact plants are seemingly corrected by light/dark cycles (Muranaka and Oyama, 2016), indicating that these cycles are extremely important to generate systemic plants responses to environmental variations. Shifts in photoperiod responsiveness were obtained over the tomato domestication process, whereas pivotal genes associated with this process have their expression intimately dependent on light/dark cycles (Xiang et al., 2022). Wild tomatoes have much more cells than domesticated tomatoes due to higher branching levels, and they are classical examples that constant light does not greatly disrupt their phenotype, in contrast to the situation in domesticated tomatoes (Velez-Ramirez et al., 2014), suggesting the synchronization of clocks in wild species. In general, different organs of wild species display variations in circadian rhythms, highlighting differences observed in cotyledon-to-cotyledon, leaf-to-leaf and shoot-to-root relations which extend to the control of the organ development (Thian, 2000; Kim et al., 2016). This may ultimately contribute to a biological advantage. It seems reasonable to anticipate therefore that, when submitted to stressful conditions, the wild species most likely suffer the loss of leaves with a lower commitment to the overall plant *fitness*. A very interesting example of synchronization may be observed in wild species *Echinacea angustifolia* which is perennial and displays a flowering time that is synchronized by fire (Wagenius et al., 2020). In domesticated tomatoes, the biological synchronization seems to have been generated due to human activity, ensuring the coordination between circadian rhythms and leaf development. It would be interesting to determine whether, and to what extent, environmental factors over the evolution of domestication may have boosted inter-organ relationships in plants.

The results obtained here also point to the developmental temporisation of domesticated tomatoes, which homogenize photosynthesis across leaves and establish stronger source-to-sink relationships. By contrast, *S. pennellii* seems to show an atemporal regulation of traits programming source-to-sink relationships, with each leaf working rather independently. It can contribute to explaining the slow growth and reduced fruit mass of *S. pennellii*, which exhibits a positive relationship with its higher ability to tolerate environmental adversities. Notably, metabolism and organism size were largely uncoupled by plant domestication (Milla et al., 2018). For instance, the highly branched pattern of wild tomatoes might be due to weak inter-tissue correlations of the metabolism, which was likely synchronized over tomato domestication. From the findings of Milla *et al*. (2018), it is possible to note that wild relatives of crops likely display a broad spectrum of relationships for metabolism *versus* plant size, suggesting that domestication indeed set developmental, circadian, and metabolic clocks in cultivated plants. Similarly, the elevated diurnal expression of *TOC1* leads ultimately to a higher starch turnover, improving biomass gain in *A. thaliana* polyploids when compared with their parental diploids (Ni et al., 2009). Consistently, *TOC1* expression and starch levels demonstrated similar patterns in domesticated tomatoes, whilst higher *TOC1* expression in L4 than L3 upon noon seems to be correlated with higher starch turnover and superior biomass gain of the young leaflet in wild relatives. In agreement, the circadian clock is particularly sensitive to sucrose in the darkness, being that *GI* plays the major role in sucrose sensing mediating shifts in rhythmicity of shoot circadian (Dalchau et al., 2011). Thus, sucrose is likely able to reset the circadian clock based on metabolic *status* acting as both signal and metabolite at the same time (Müller et al., 2014). *GI* expression was highly homogenous between L3 and L4 whereas sucrose levels were widely divergent for both leaflets over the diel period in M82, indicating that sucrose is not able to reset rhythms of L3 and L4. The opposite was observed for *S. pennellii* that showed a contrasting *GI* expression between L3 and L4, despite almost similar sucrose levels for both leaflets. It seems, therefore, that the contrasting development, physiology, and metabolism between older and younger leaves of *S. lycopersicum* likely program the source-sink relationships, which seems to have been reached during its domestication.

Over the past decades our understanding of circadian clocks, development and metabolism experienced significant advances, wherein the function of thousands of genes, proteins, and metabolites have been demonstrated (Greenham and McClung, 2015; Yu et al., 2015). Notwithstanding this fact, the structure of circadian, developmental, and metabolic clocks along plant species evolution is far from being fully understood. We posit that distinct plant tissues may exhibit variations in inter-relationships over plant development. Domesticated tomato is an interesting case that may contribute to starting to solve how biological clocks are structured in plants, while *de novo* domestication provides an outstanding opportunity to re-structure these clocks to engineer ideal crops. The results described here provide compelling evidence for a higher coherence among tissues from domesticated tomato, which seemingly does not occur in their wild relatives. Future studies must explore the temporal (in)coherences among organs and tissues to generate ideal crops that are more productive under fluctuating environments, ensuring food security even in hostile environments.

## Data availability

The data that support the findings of this study are available from the corresponding authors upon reasonable request.

## Acknowledgments

Discussions with Professor Wagner Otoni (Universidade Federal de Viçoa, Brazil), Lázaro E. P. Peres (Universidade de Sao Paulo, Brazil), Cristiane Calixto Universidade de Sao Paulo, Brazil) were highly valuable for the development of this work. This work was supported by the Serrapilheira Institute-Brazil [grant number: Serra-1812-27067 to W.L.A.], National Council for Scientific and Technological Development (CNPq-Brazil), and Foundation for Research Assistance of the Minas Gerais State, (FAPEMIG-Brazil). We also acknowledge research fellowships granted by CNPq to A.N-N. and W.L.A.

## Author Contributions

J.A.S., and W.L.A. designed the research; J.A.S. performed most of the research with the support of A.O.M., T.W., M.F.S., W.B-S. and F.L.; A.R.F., and A.N-N. contributed new reagents/analytic tools; J.A.S., A.R.F., A.N-N., and W.L.A. analyzed the data; and J.A.S., and W.L.A. wrote the article with input from all the others.

