## Supplemental Figures for "The temporal regulation inter-leaves from domesticated-tomato contrasts with timelessness of its wild ancestors"

### Supplementary Figures

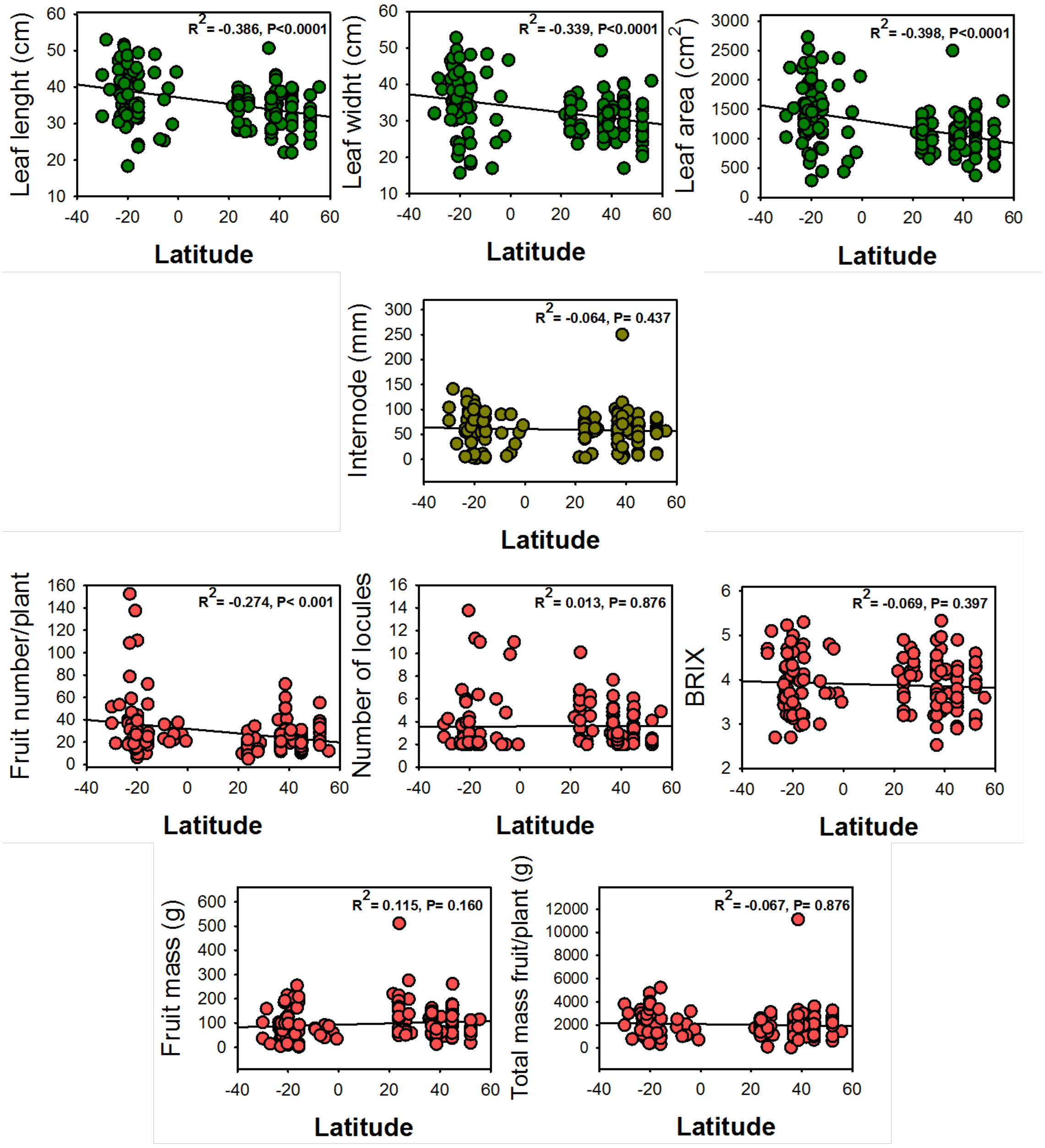


**Supplementary figure S1. Pearson correlations for vegetative and reproductive traits of *Solanum lycopersicum* accessions.** The data were obtained from germplasm bank at the Universidade Federal de Viçosa, Brazil (www.bgh.ufv.br). Values are presented as correlation index ($R^2$) and *p-value*.

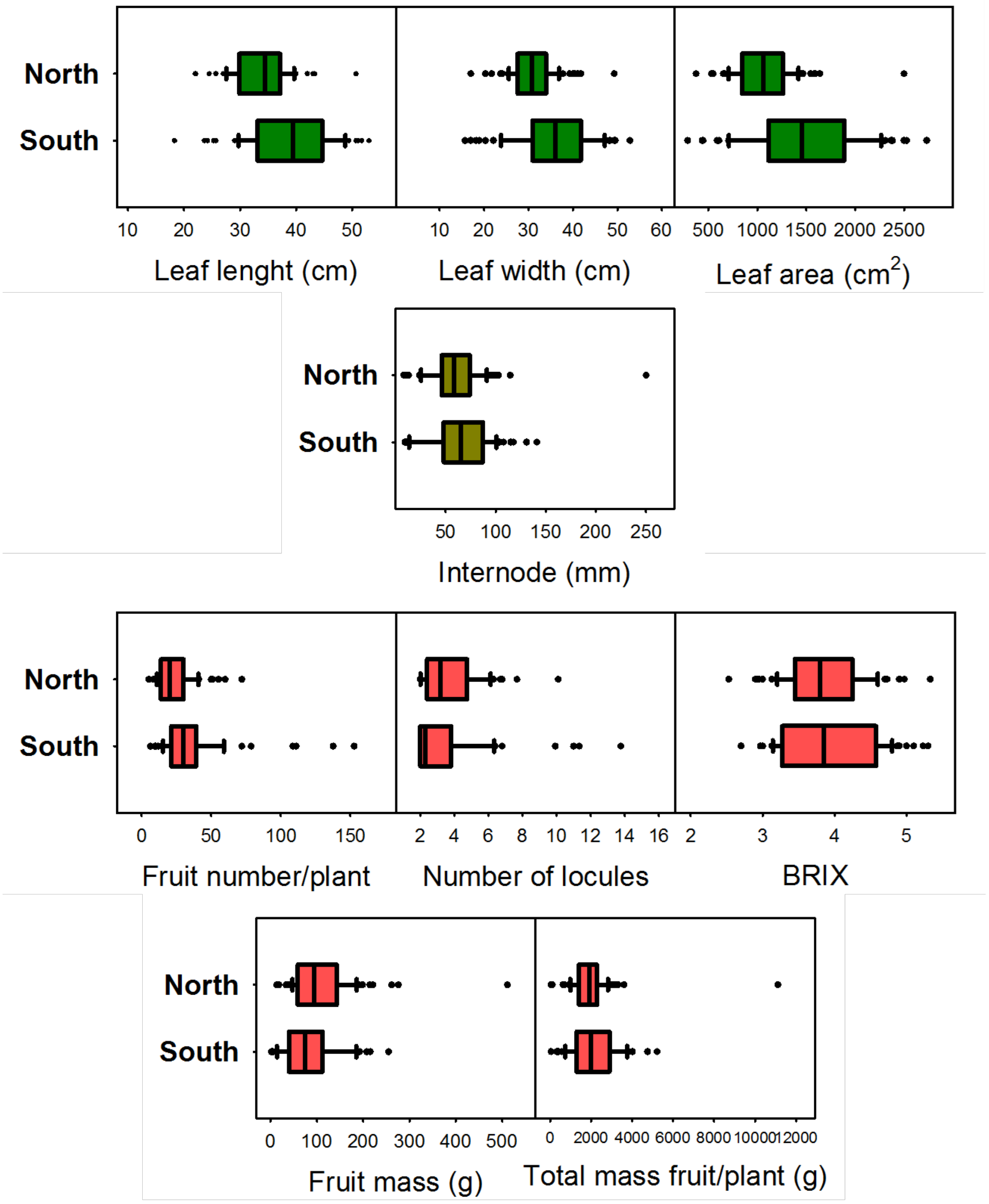


**Supplementary figure S2. Comparisons for vegetative and reproductive traits of *Solanum lycopersicum* accessions originated from the north and south hemispheres.** The data were obtained from germplasm bank at the Universidade Federal de Viçosa, Brazil (www.bgh.ufv.br/).

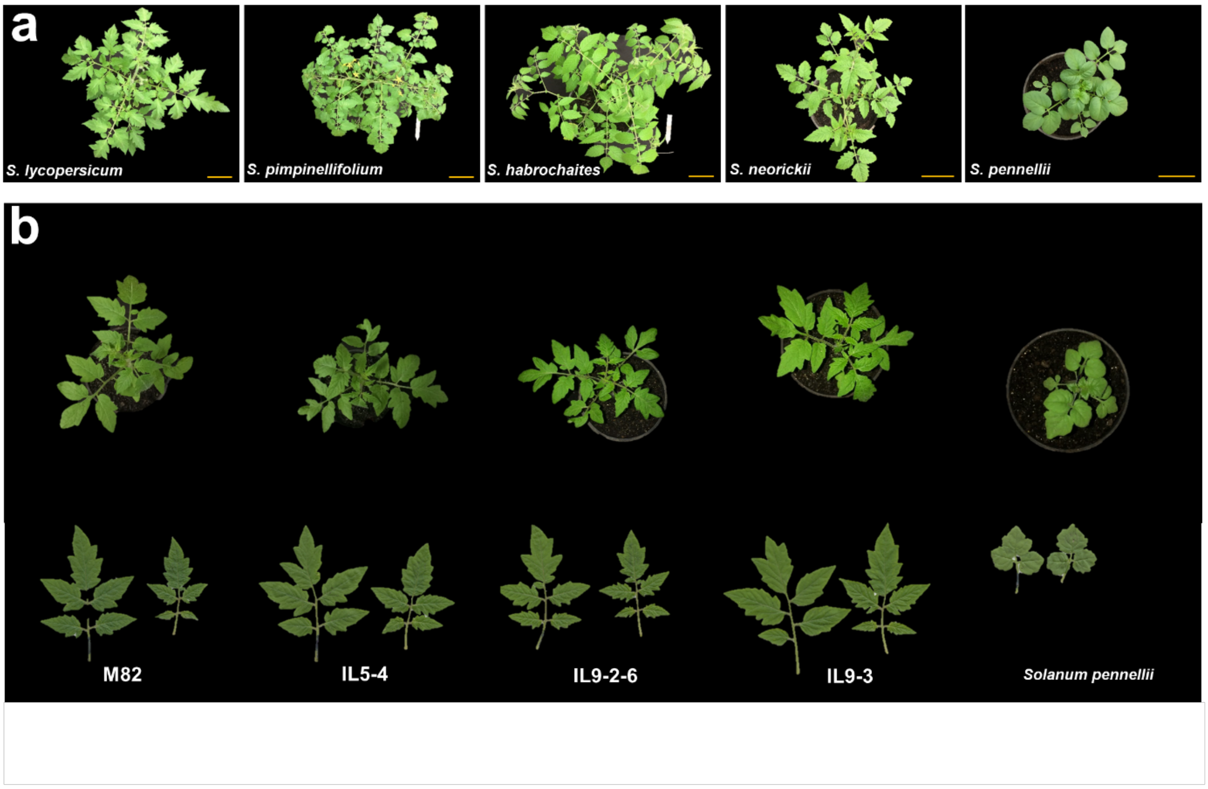


**Supplementary Figure S3. Phenotypical coherence on tomato development. a,** branching patterns (octave week) across *Solanum* species, where it showed the domesticated tomato *Solanum lycopersicum* (cv. M82) and the wild tomatoes *Solanum pennellii*, *Solanum neorickii*, *Solanum habrochaites,* and *Solanum pimpinellifolium*also. **b,** branching patterns and phenotype for the 3$^{rd}$ and 4$^{th}$ leaves at fourth week, the image represents cv. M82 and *S. pennellii* as well as introgression lines (IL) on M82 containing genomic fragments of *S. pennellii*.

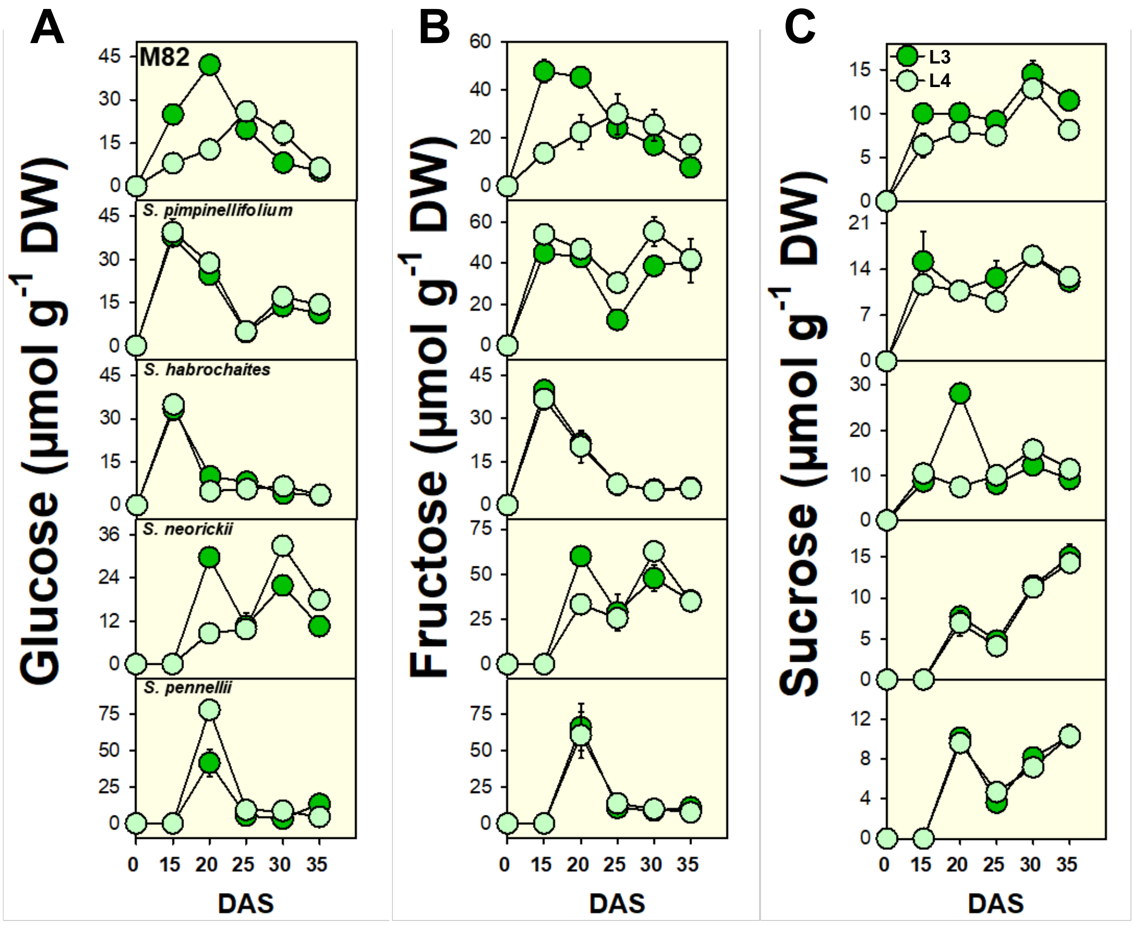


**Supplementary Figure S4. The metabolism over leaf development on wild and domesticated tomatoes.** Older leaflets of third (dark green) and fourth (pale-green) leaves from domesticated tomato (*Solanum lycopersicum*, Cv. M82), *S. pimpinellifolium*, *S. habrochaites*, *S. neorickii*, and *S. pennellii*. It is represented the levels of glucose, fructose, and sucrose. Sugar levels were assessed in leaflets harvested every five days over development. Data are means ± SE (n = 5). DW: Dry weight; DAS: Days after sowing.

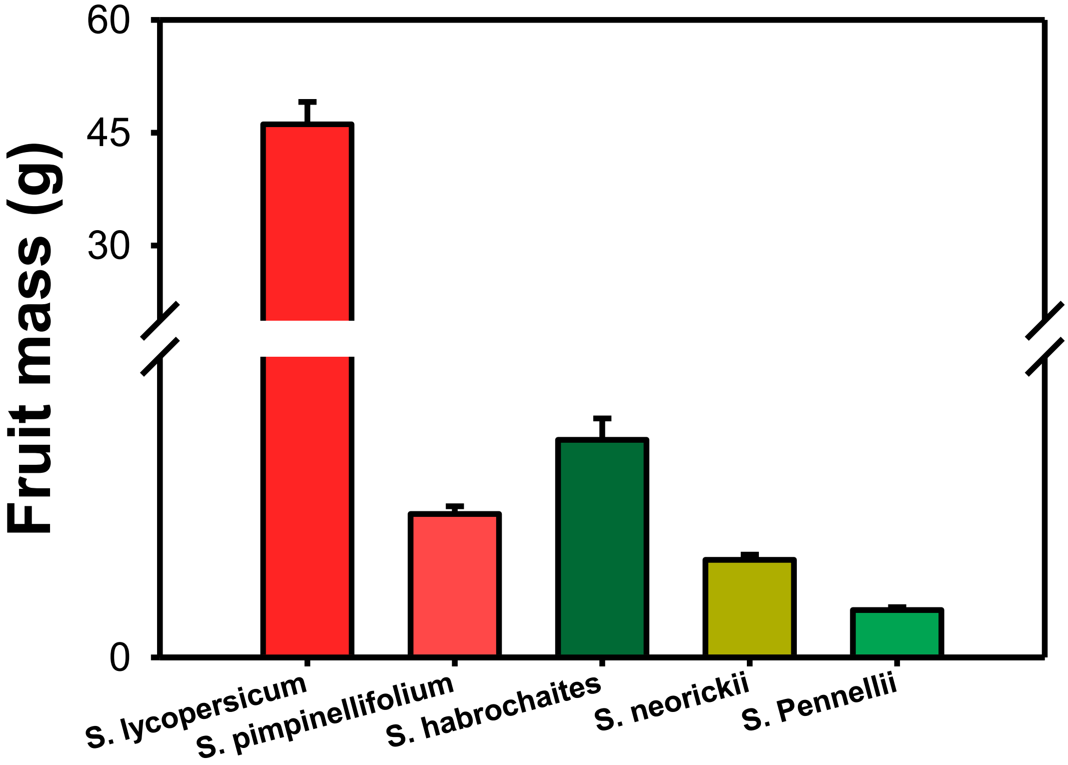


**Supplementary figure S5. Fruit mass for tomato species (g).** The fruits were harvested over plant development. Values are presented as means ± SE of individual fruits (n = 5 individual plants).

| Trait* | M82 | | IL5-4 | | IL9-2-6 | | IL9-3 | | *S. pennellii* | |
|---|---|---|---|---|---|---|---|---|---|---|
| **Leaf** | 3rd | 4th | 3rd | 4th | 3rd | 4th | 3rd | 4th | 3rd | 4th |
| LA | 23.9±2.4 | **11.9±1.5** | 21.5±2.2 | 15.4±2.1 | 18.7±1.9 | 15.6±1.5 | 25.9±2.9 | 15.6±1.5 | 5.4±0.5 | 5.1±0.7 |
| LL | 10.2±0.4 | **7.7±0.4** | 9.7±0.5 | 8.3±0.4 | 9.45±0.5 | 8.4±0.5 | 10.3±0.7 | 9.4±1.1 | 3.4±0.4 | 3.3±0.3 |
| LM | 34.1±3.9 | **21.7±2.5** | 35.4±3.5 | 26.0±3.3 | 29.11±2.9 | 22.6±2.8 | 34.0±4.9 | 27.7±4.7 | 6.2±1.0 | 6.5±0.9 |
| SLA | 671±17 | **601±14** | 639±9 | 627±14 | 680±11 | **630±5** | 647±49 | 714±68 | 770±38 | 714±36 |

**Supplementary Table 1. Leaf growth traits on *Solanum lycopers*icum (cv. M82), *Solanum pennellii* and on introgression lines (IL).** Data presented are mean ± SE (n = 7) obtained in at least two independent assays. Bold numbers describe differences $P < 0.05$ between 3rd and 4th leaves for each genotype, which it was calculated by two-sided Student's *t*-test.

* LA: leaf area ($cm^2$); LL: leaf length (cm); LM: leaf mass; SLA: specific leaf area ($cm^2$ $g^{-1}$).

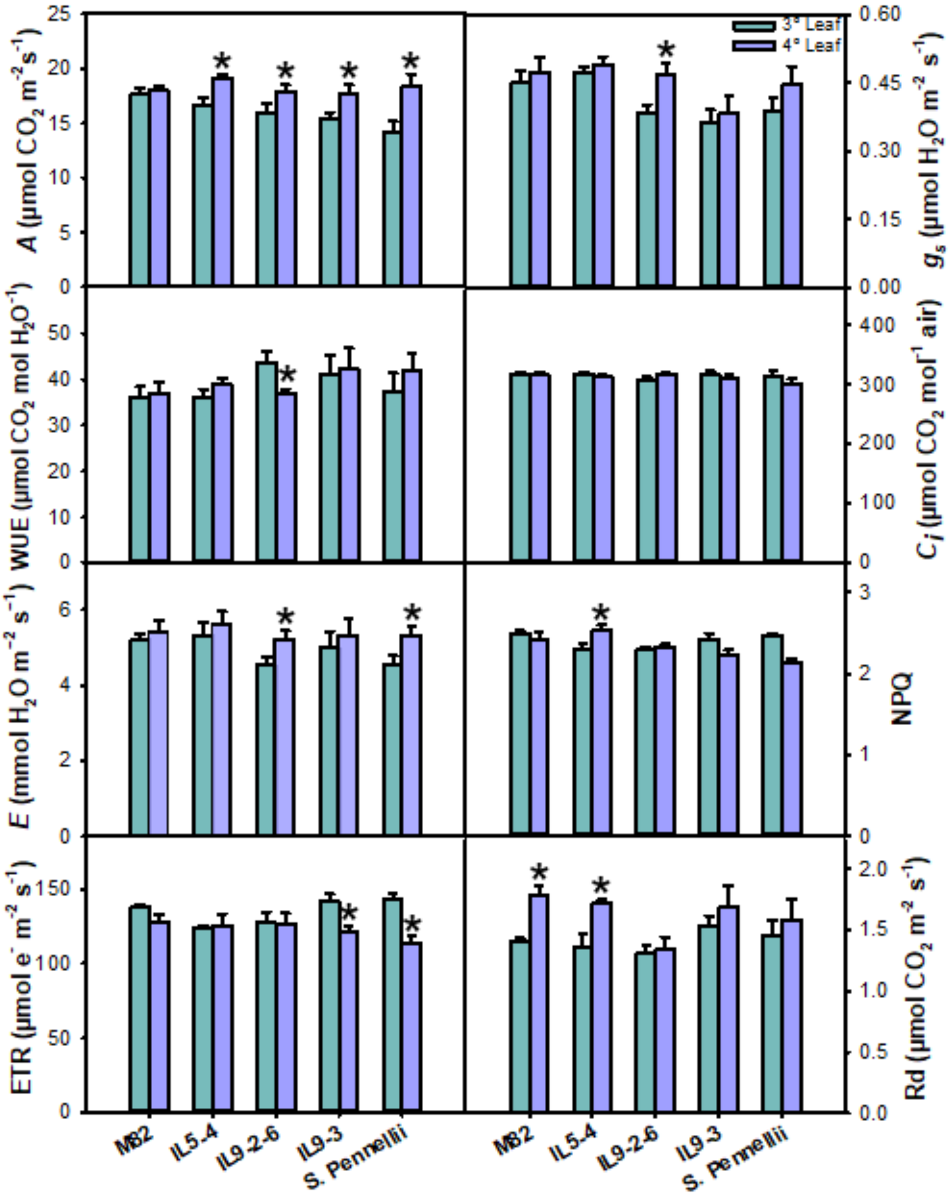


**Supplementary Figure S6. Gas exchange parameters exhibit a differential regulation.** Data were obtained in 4-week-old plants Solanum lycopersicum (cv. M82), Solanum pennellii and on introgression lines (IL) growing under optimal conditions. Values are presented as means ± SE (n = 5). Asterisks denotes differences ($P < 0.05$) between the 3$^{rd}$ and 4$^{th}$ leaves for each genotype, which it was calculated by two-sided Student's *t*-test. *A*: photosynthesis rate; $g_s$: stomatal conductance; WUE: intrinsic water-use efficiency; $C_i$: internal $CO_2$ concentration; E: transpiration rate; NPQ: non-photochemical quenching; ETR: electron transport rate; $R_d$: dark respiration.

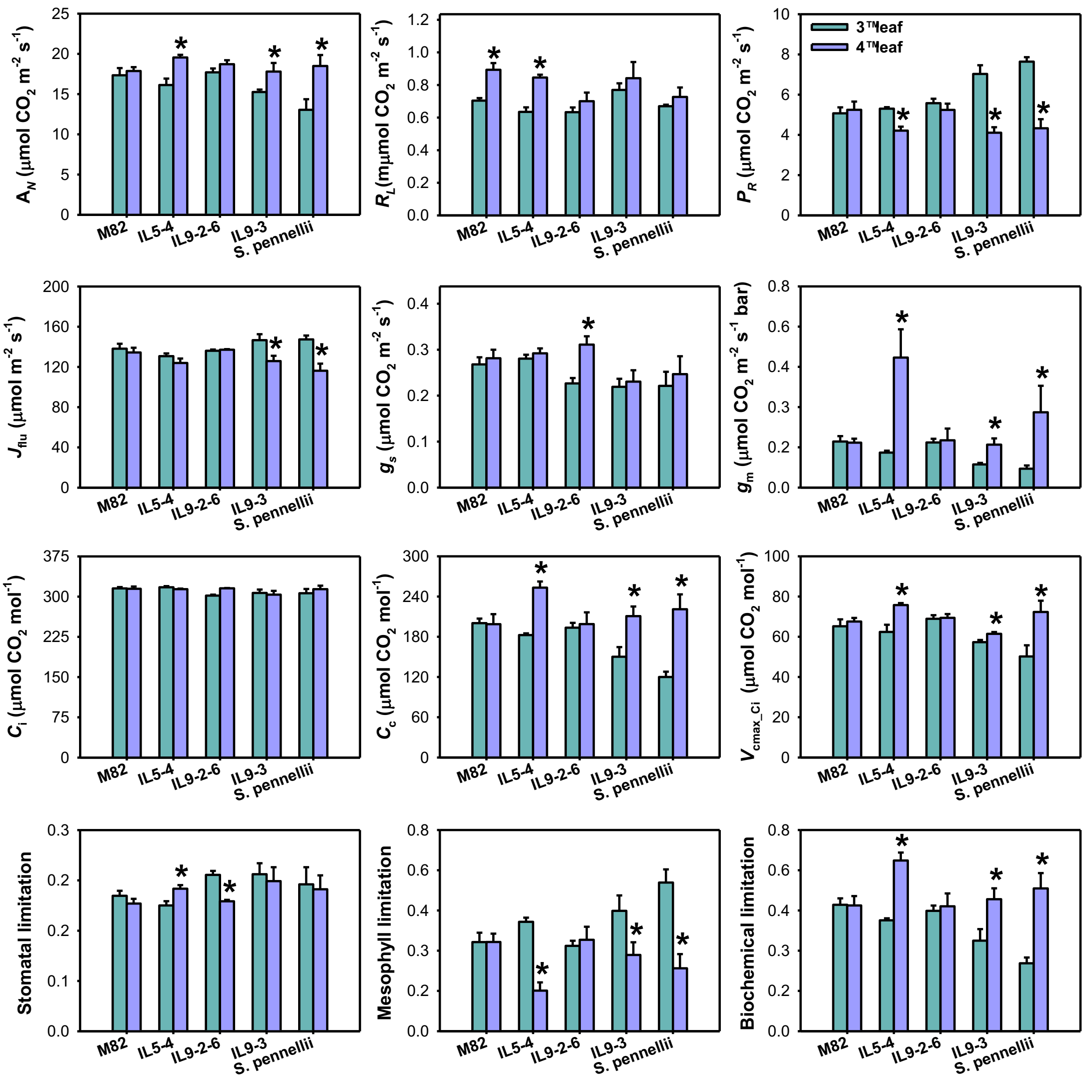


**Supplementary Figure S7. Influence of tomato domestication on photosynthetic efficiency of different plant leaves.** Estimative for photosynthesis traits, where the data were obtained in 4-week-old plants *Solanum lycopersicum* (cv. M82), *Solanum pennellii* and on introgression lines (IL) growing under optimal conditions. Values are presented as means ± SE (n = 5). Asterisks denotes differences ($P < 0.05$) between the 3$^{rd}$ and 4$^{th}$ leaves for each genotype, which it was calculated by two-sided Student's *t*-test. $A_N$: Net photosynthesis rate; $R_L$: light respiration; $P_R$: photorespiration rate; $J_{flu}$: electron transport rate estimated by chlorophyll fluorescence parameters; $g_s$: stomatal conductance $CO_2$; $g_m$: mesophyll conductance; $C_i$: internal $CO_2$ concentration; Cc: chloroplast $CO_2$ concentration; $V_{cmax\_Ci}$: Rubisco maximum carboxylation capacity based on $C_i$.

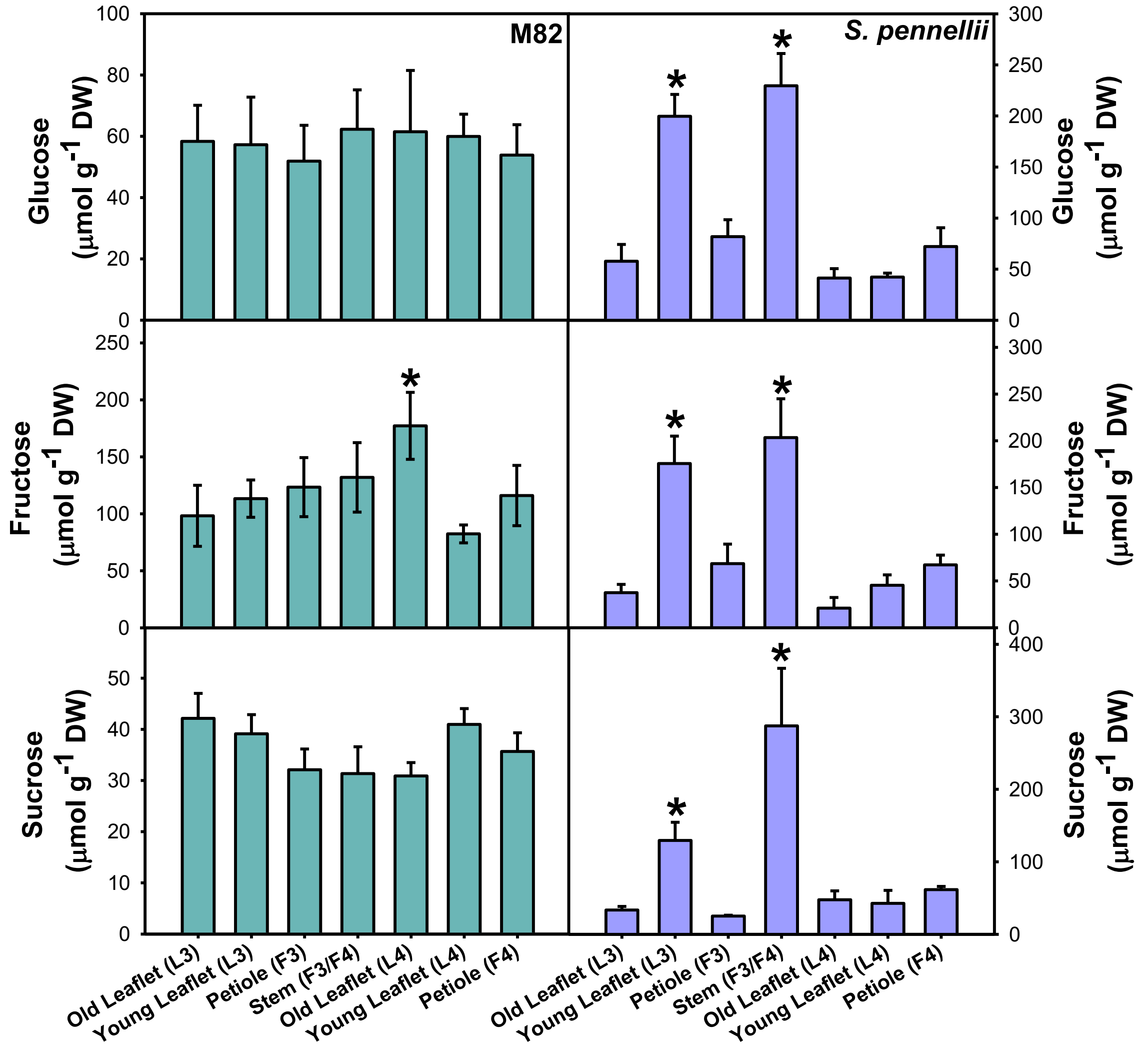


**Supplementary Figure S8. Leaf metabolite levels are altered on tomato segments**. Key metabolite content in leaves of tomato plants: glucose, fructose, and sucrose in samples harvested at the middle of the day. Samples were taken from the third and fourth leaf from the apex of 4-week-old plants. Data are means ± SE (n = 5), and asterisks describe differences ($P$ < 0.05) between the 3$^{rd}$ (L3) and 4$^{th}$ (L4) leaves for each genotype, which was calculated by two-sided Student's $t$-test. DW: Dry weight.
